# Structural basis of ATP release and ion selectivity in Pannexin1 channels

**DOI:** 10.1101/2025.02.17.638584

**Authors:** Qingyang Zhang, Junjie Wang, Guillaume Gaullier, David Kotol, Cecilia Blikstad, Gerhard Dahl, Rene Barro-Soria, Carsten Mim

## Abstract

This study investigated the structural basis of adenosine triphosphate (ATP) release and ion selectivity by Pannexin1 (Panx1). The microinjection of active SRC tyrosine kinase increased Panx1-mediated currents in Panx1-expressing oocytes. Furthermore, a mutation mimicking phosphorylation at Y308 turned Panx1 into a constitutively open ATP-releasing channel with broad ion permeability. Cryogenic electron microscopy showed that for most maps, the channel alternated between wide and narrow conformations. However, in the map of the phosphomimetic Y308E mutation, Panx1 remains in a wide ATP-releasing conformation. The pore-lining residues, W74 and R75, are flexible and create, in concert with the sequestered N-terminal helix, a path for ATP. The data point to phosphorylation as a reversible switch that activates ATP permeability by stabilizing Panx1 in the ATP-releasing conformation, which is crucial for developing modulators for diseases such as cancer and chronic pain.

## Introduction

Pannexin1 (Panx1) forms membrane channels that are central to physiological and pathophysiological processes, including neuronal maturation, synaptic plasticity, inflammasome activation, and neutrophil/macrophage chemotaxis in the central and peripheral nervous systems (Adamson and Leitinger, 2014; Lemaire et al., 2011), stroke (Navis et al., 2020; Thompson et al., 2006), chronic pain (McAllister et al., 2024; Yeung et al., 2020), and cancer (Kepp et al., 2021; Varela-Vazquez et al., 2020; Vultaggio-Poma et al., 2020). Depending on the stimulus quality, Panx1 channels either adopt a “large” or “small” pore conformation. The “large” pore conformation allows for the extracellular release of large signaling molecules, including adenosine triphosphate (ATP), and the passage of positively charged tracer molecules such as YoPro-1, 4′,6-diamidino-2-phenylindole, or propidium iodide (Bao et al., 2004; Chekeni et al., 2010; Dahl, 2015; Garre et al., 2010; Qiu et al., 2012; Wang and Dahl, 2018). Conversely, the “small” pore conformation of Panx1 serves as a chloride-selective ion channel (Dahl, 2015; Wang et al., 2014; Wang and Dahl, 2018) and has no cation and ATP permeability (Ma et al., 2012; Romanov et al., 2012; Wang et al., 2014). The “small,” chloride-selective pore conformation is stimulated by membrane depolarization or proteolytic cleavage of the carboxy terminus of Panx1 by caspases. The “large,” ATP-permeable pore conformation is either directly stimulated by mechanical stress and low oxygen levels (e.g., in erythrocytes) or indirectly through membrane receptors (Bao et al., 2004; Locovei et al., 2006b; Murali et al., 2014; Thompson et al., 2006; Weilinger et al., 2012).

The proposed proteolytic gating mechanism (Chekeni et al., 2010; Narahari et al., 2021; Ruan et al., 2020; Sandilos et al., 2012) activates apoptosis and does not align with the biological necessity of reversible activation. However, the reversible activation of the “large” pore conformation of Panx1 by phosphorylation through SRC tyrosine kinase is a distinct possibility. This was first described in the context of the interaction between Panx1 and the P2X7 receptor (Iglesias et al., 2008). Other receptors, including N-methyl-D-aspartate receptors (NMDARs), activate Panx1 via SRC phosphorylation (Leonard et al., 2018; Leonard et al., 2019; Weilinger et al., 2016). In mice, SRC targets at least four tyrosine residues: Y10, Y150, Y198, and Y308 (DeLalio et al., 2019; Langlois et al., 2023; Lohman et al., 2019; Metz and Elvers, 2022; Nouri-Nejad et al., 2021; Weilinger et al., 2016; Weilinger et al., 2012). However, the mechanism by which Panx1 is activated through the phosphorylation of these tyrosine residues is poorly understood. The phosphorylation of Y10 and Y150 may affect trafficking by altering glycosylation (Langlois et al., 2023; Nouri-Nejad et al., 2021). Tyrosine Y198 has been shown to be constitutively phosphorylated and, consequently, cannot serve as a switch for conformational change into the “large” pore conformation of Panx1 (DeLalio et al., 2019). Conversely, tyrosine Y308 phosphorylation could serve as such an activation switch (Weilinger et al., 2016). A recent study rejected the idea that human Panx1 is phosphorylated at Y199 (Y198 in mice) and Y309 (Y308 in mice) based on negative results obtained using established phosphorylation-sensitive Panx1 antibodies and mass spectrometry (Ruan et al., 2024). Therefore, the phosphorylation of Panx1 and the potential regulation of Panx1 channel activity require further functional and structural elucidation.

The cryogenic electron microscopy (cryo-EM) structures and models of Panx1 show that tryptophan at position 74 (W74) creates the narrowest constriction at the extracellular pore with a maximum diameter of approximately 9 Å. Studies have suggested rigid positioning of residue W74 and the adjacent arginine (R75) (Michalski et al., 2020; Qu et al., 2020; Ruan et al., 2020). This assembly is stabilized via a strong cation-π interaction between R75 and W74 (Infield et al., 2021). The R75-W74 pair is also essential for ion selectivity, the ATP feedback mechanism, and possibly ATP release (Hussain et al., 2024; Michalski et al., 2020; Qiu et al., 2012; Qu et al., 2020). However, the latter function is contradicted by functional data from the alanine substitution mutant W74A, which does not result in an ATP-releasing channel without stimulation (Qiu et al., 2012). In addition, while all current structure-function studies support the “small” chloride-selective conformation, the “large” pore conformation remains elusive (Mim et al., 2021; Syrjanen et al., 2021).

Therefore, this study aimed to investigate the structural basis of ATP release and ion selectivity by Panx1, focusing on understanding how phosphorylation affects the function and conformation of Panx1. To this end, we obtained the cryo-EM structures of *in vitro*-phosphorylated (Panx1SRC) and dephosphorylated (Panx1Deph) mouse Panx1, a phosphomimetic Y308E mutant, and the pore-lining alanine mutants R75A and W74A. In addition, we characterized Y308E and *in vitro*-phosphorylated Panx1 electrophysiologically. Our findings suggest that phosphorylation is the long-sought reversible physiological switch for ATP permeability in Panx1. We propose that phosphorylation alters the protein dynamics of Panx1 in favor of a wide state. This result is critical to developing Panx1 modulators for diseases such as cancer and chronic pain.

## Results

### v-SRC increases Panx1 current and induces structural flexibility

Previous reports have suggested that the activation of NMDAR and other receptors (Iglesias et al., 2008; Leonard et al., 2019; Weilinger et al., 2016) induces the “large” pore conformation of Panx1 via SRC-mediated Panx1 phosphorylation. Diverging reports (DeLalio et al., 2019; Ruan et al., 2024) motivated us to test whether SRC alters Panx1-mediated currents. We recorded currents using two-electrode voltage clamp (TEVC) electrophysiology in mouse-Panx1-expressing *Xenopus* oocytes while injecting them with a constitutively active form of SRC (v-SRC). A single v-SRC injection induced Panx1-mediated outward currents in excess over those following albumin injection. Injection of active v-SRC into control oocytes (preinjected with water rather than mRNA) remained without noticeable effects (Fig. 1A, B). Using mass spectrometry, we mapped possible v-SRC phosphorylation sites *in vitro* using purified mouse-Panx1 (hereafter referred to as Panx1). Detergent-purified Panx1 was dephosphorylated with λ-phosphatase and rephosphorylated with v-SRC *in vitro*. We identified only one Panx1 tyrosine phosphorylation site (Y308) that was significantly enriched following v-SRC treatment. We identified two phosphoserine residues, S181 and S406 (Fig. 1C). Y10 was phosphorylated but not significantly enriched compared with the dephosphorylated sample (Fig. S9B).

**Fig. 1.**
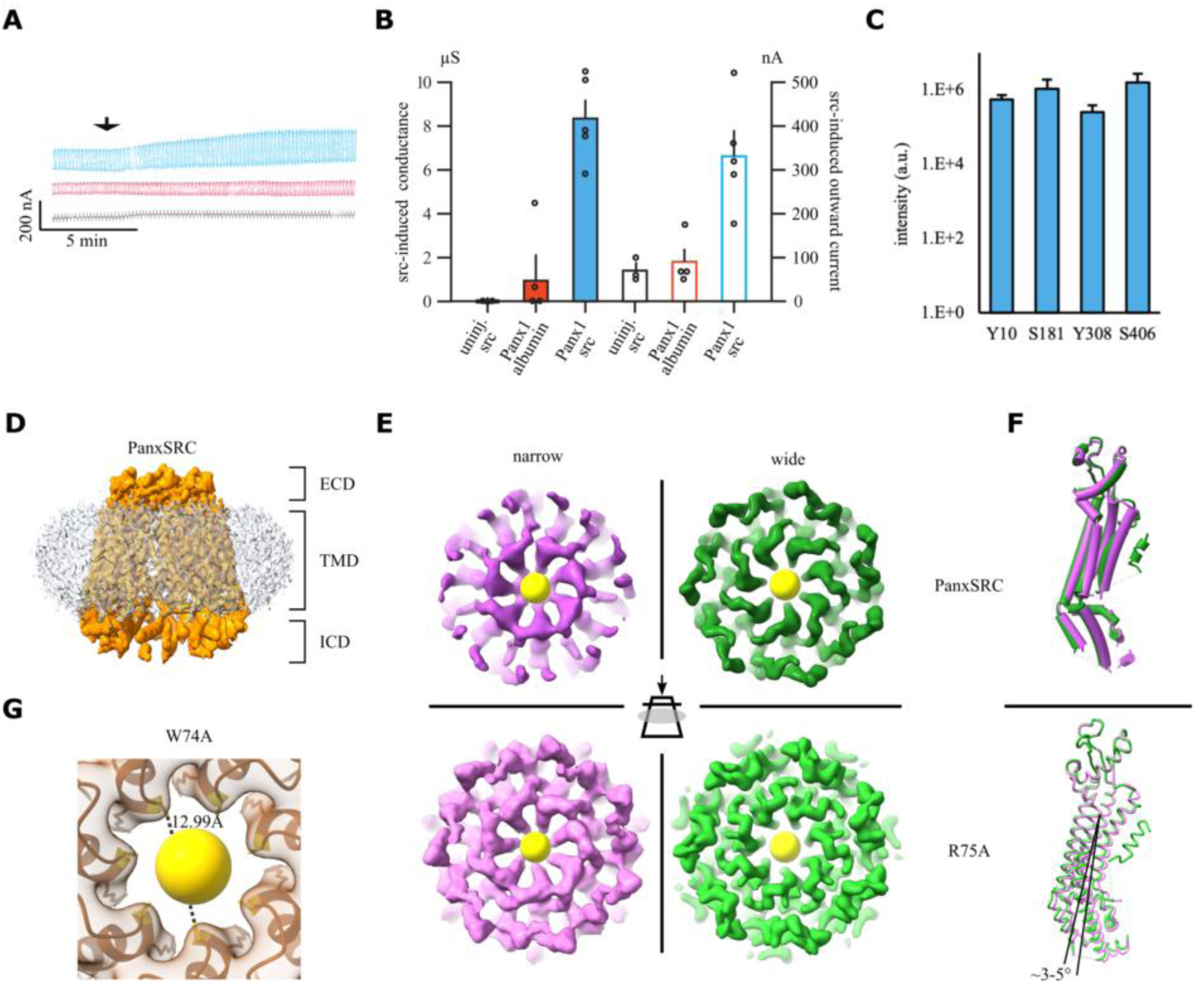
Characterization of v-SRC dependent phosphorylation of Panx1 and its conformation dynamics. (**A**) Current traces of oocytes clamped at –60 and 20 mV depolarizing test pulses at 0.1 Hz. Arrowheads indicate the time of v-SRC injection. Blue trace: injection of 37 nL of active v-SRC protein increased the Panx1-mediated currents. Red trace: injection of the same volume of albumin solution (10 µg/mL) remained without effect. Black trace: injection of active v-SRC into native control oocytes did not affect endogenous channels. (**B**) Quantitative analysis of v-SRC-induced conductance changes (filled bars) and v-SRC-induced currents (empty bars) for Panx1 mRNA uninjected oocytes (black, *n* = 3), albumin injected into Panx1-expressing oocytes (red, *n* = 4), and v-SRC injected into Panx1-expressing oocytes (blue, *n* = 5). All error = SD (**C**) Intensity of phosphorylated peptides at four sites in the v-SRC treated cryogenic electron microscopy (cryo-EM) sample (*n* = 3, error = SD). (**D**) Sharpened cryo-EM map of Panx1 phosphorylated with v-SRC (PanxSRC, orange) viewed in the plane of the membrane with the detergent micelle (gray) extracellular domain (ECD), transmembrane domain and intracellular domain labeled. (**E**) Volumes from three-dimensional variability analysis (3DVA) of PanxSRC (top) and R75A (bottom) particles. The view is an extracellular view through the plane in the ECD, indicated by the illustration in the center. The narrow conformation is colored in purple, and the wide conformation is in green. A yellow sphere was placed at the center of the pore for illustration. (diameter 8 Å). (**F**) Overlay of the atomic models refined against the first and last frames of the 3DVA for one Panx1 protomer: PanxSRC (pipes and planks) and R75A (licorice). The view is in the plane of the membrane, as in (**D**). The wide form (green) sees a shift in the transmembrane domain (TMD) compared with the narrow form (purple). (**G**) View from the cytosol to the extracellular space for the sharpened cryo-EM map of W74A (transparent brown) with its model (brown ribbon). The distance between opposing residues is 12.9 Å. A sphere was placed in the center of the pore for orientation (diameter 12 Å).

Generally, the presence of non-phosphorylated peptides at these four sites (Y10, S181, Y308, and S406), particularly Y308, in the v-SRC-treated samples indicated that some Panx1 subunits or whole Panx1 channels were not phosphorylated (Fig. S9C). Using cryo-EM, we assessed the structural impact of the two analyzed samples, namely, the dephosphorylated form of Panx1 (PanxDeph) and PanxDeph treated with v-SRC (PanxSRC). PanxSRC yielded a map with a resolution of 3.6 Å (Fig. 1D and S1.3), and PanxDeph resolved to 4.7 Å (Fig. S1.5).

The overall fold and domain organization of PanxDeph and PanxSRC were similar to those of other Panx1 structures (reviewed in (Syrjanen et al., 2021)). The channel formed heptamers, each comprising four transmembrane helices and a pore constricted by tryptophan at position 74 (W74). However, similar to other cryo-EM maps (e.g., (Drulyte et al., 2023; Hussain et al., 2024; Qu et al., 2020; Ruan et al., 2020)), we observed a steady decrease in local resolution from the extracellular domain (ECD) to the intracellular domain (ICD). This suggested conformational heterogeneity, and we therefore analyzed the particle set using three-dimensional variability analysis (3DVA) (Punjani and Fleet, 2021). Our analysis revealed that PanxSRC exists in narrow-(Fig. 1E, top purple) and wide-pore conformations (Fig. 1E, top green; Movie S1). After further processing (Movie S2), we separated and reconstructed the particles associated with the most extreme maps and built models (Fig. 1E, purple vs. green = maps, Fig. 1F = model). The hallmarks of the transition between both conformations are the presence or absence of density for the N-terminal helix (NTH), unfolding of the C-terminal end of helix S1, and tilting of the transmembrane helices (Fig. 1F, Fig. 4D, Fig. S8B, see S10 for naming of structure elements). The narrow-pore model shares similarities with the PanxDeph map, and the structure of probenecid-inhibited Panx1 (Kuzuya et al., 2022).

To investigate the role of the pore-lining residues, we determined the structures of the R75A and W74A mutants (Fig. 1E, lower panels, Fig. 1G) resulting in an overall resolution of 2.9 and 3.8 Å, respectively (Fig. 1E, bottom panels, Movie S3, and Fig. 1G). Additionally, the overall fold was similar to that of previous structures (Mim et al., 2021; Syrjanen et al., 2021). Visualization of the conformational dynamics provided valuable insights into PanxSRC. Therefore, we also performed 3DVA for W74A and R75A. The W74A (Movie S6) and R75A (Movies S3 and S4) mutants displayed structural dynamics similar to that of PanxSRC with respect to the movement of the transmembrane domain (TMD; Fig. 1F, bottom for R75A). Notably, we observed that the density in the pore varied between the narrow and wide conformations, implying movement of W74 in the pore during the transition (Fig. 1E, Movies S1-4). The W74A mutant generated channels whose pore (approximately 12 Å in diameter) was wide enough to allow ATP permeation (Fig. 1G and Fig. S1.4). This result suggests that fully retrieving the bulky W74 from the narrowest point of the permeation pathway (e.g., swinging W74 away) may widen the pore sufficiently to accommodate large molecules such as ATP. Despite diverging results, functional data suggest a more complex mechanism because W74A may not be an ATP-releasing channel and requires extracellular K^+^ stimulation for ATP release (Qiu et al., 2012) vs. (Qu et al., 2020).

### Phosphomimetic mutants create a constitutively ATP-permeable channel with altered ion selectivity

The Panx1-W74A mutant might not release ATP unstimulated (Qiu et al., 2012), indicating that phosphorylation has an additional effect beyond moving W74. We also showed that v-SRC may not phosphorylate all sites in the heptameric Panx1 channel in our sample (Fig. S9C). Therefore, we designed mutations that mimic phosphorylation (phosphomimetic), an approach commonly used to investigate ion channel function (Barvikova et al., 2020). Based on our data and previous reports on functionally important sites (DeLalio et al., 2019), we introduced phosphomimetic (Y10E, Y198E, and Y308E) and phosphosuppressive (Y198F) single and double mutants (Fig. 2A).

**Fig. 2.**
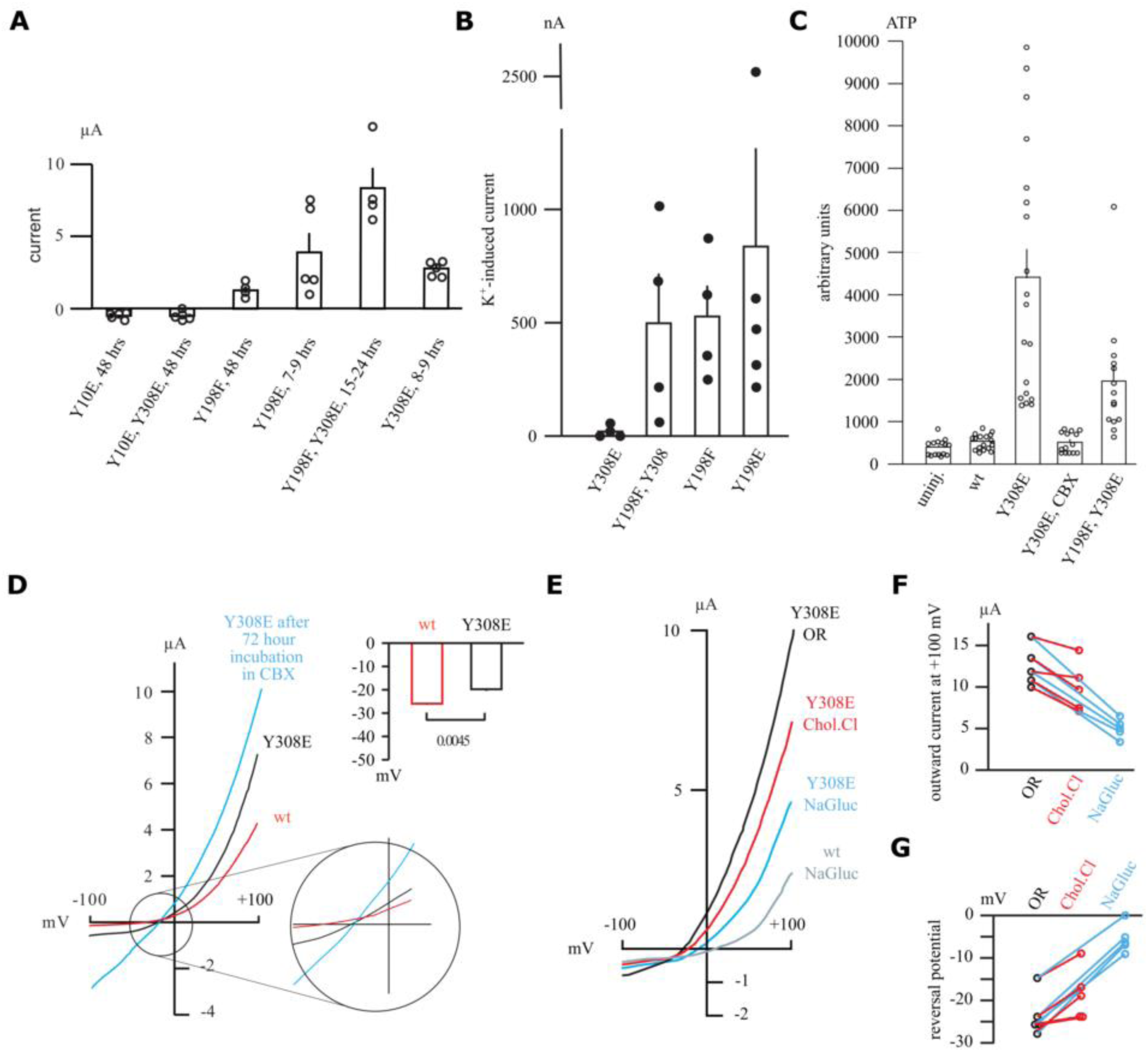
Functional characterization of Panx1 phosphomimetic mutants. (**A**) Membrane currents for oocytes expressing Panx1 phospho-site mutants. Panx1Y10E (*n* = 5) and Panx1Y10E/Y308E (*n* = 5) exhibit small inward currents. Similar inward currents are observed in mRNA uninjected oocytes and are likely endogenous sodium channels (Baud et al., 1982; Methfessel et al., 1986). Y198F (*n* = 4), Y198E (*n* = 5), Y198F, Y308E (*n* = 4), and Y308E (*n* = 5) exhibited CBX-sensitive, outward currents with different expression kinetics. (**B**) Oocytes expressing mutants Y198F/Y308E, Y198E, and Y198F exhibited inward currents and increased membrane conductance when voltage clamped at –60 mV and perfused with 85 mM [K^+^]. In contrast, Y308E could not further be stimulated by perfusion with 85 mM [K^+^]. (**C**) ATP release by oocytes in the absence of stimulus. mRNA uninjected oocytes (*n* = 15) and Panx1-expressing oocytes (*n* = 16) do not release ATP. Oocytes expressing Y308E alone (*n* = 19) or with a phoshosuppressive mutant (Y198F/Y308E, *n* = 14) show large ATP release. ATP release can be inhibited by the channel blocker CBX (*n* = 14). (**D**) Current traces induced by voltage ramps from –100 to +100 mV are shown for oocytes expressing wtPanx1 for 24 h (red), Panx1 Y308E for 8 h (black), or Y308E for 72 h in CBX (blue) after mRNA injection. For Y308E, CBX was washed out before measurements. The inset shows the enlarged current trace intersect (bottom) and the average reversal potential (top). (**E**) Current traces induced by voltage ramps from –100 to +100 mV recorded for oocytes expressing Y308E for 8 h after injection in the presence of different ions: black = Oocyte Ringer’s solution (OR), red = CholineCl Ringer’s solution (ChoCl), blue = NaGluconate Ringer’s solution (NaGluc). For reference, a trace of wtPanx1 expressing oocytes in gluconate solution is shown (gray wtNaGlu). Current amplitudes and reversal potentials were lowered by gluconate and choline in the OR solution. However, Y308E is less affected by the NaGlu change than wtPanx1, indicating a difference in ion selectivity. (**F**) Quantitative analysis of Y308E currents at +100 mV with the same color and naming scheme as in **E**. (**G**) Quantitative analysis of Y308E reversal potentials of the currents in the same solutions and following the same experimental and naming scheme as in **E** and **F**.

Oocytes expressing Y198E or Y308E showed outward currents as early as 7 h post-injection. Notably, cells expressing Y308E died after 24 h unless protected by the Panx1 channel blocker carbenoxolone (CBX) (Fig. S2B). Cells injected with F198F/Y308E and Y198F mRNA showed outward currents after 24 h, as typically observed for wtPanx1(Wang and Dahl, 2018) (Fig. 2A). These currents were sensitive to CBX (Fig. S2A). The phosphomimetic single and double mutants, Y10E and Y10E/Y308E, showed no outward currents (Fig. 2A, Fig. S2B). All phosphomimetic mutants, except Y10E, showed increased Panx1-mediated outward currents. We tested whether this translated into ATP release without stimulation (Fig. 2C). Cells expressing Y308E released large amounts of ATP compared with those expressing wtPanx1 channels (Fig. 2C), which CBX could block. The Y198F/Y308E double mutant, albeit less, also released ATP in the absence of a stimulus, indicating that Y198 phosphorylation is not essential for the channel to assume the “large” pore conformation (Fig. 2C).

W74 regulates ion selectivity, and alanine substitution attenuates the rectification of Panx1(Michalski et al., 2020). Therefore, we tested whether the phosphomimetic Y308E mutation changes the reversal potential of Panx1 as a means of evaluating pore changes (Fig. 2D). Oocytes expressing wtPanx1 channels (24 h after mRNA injection) exhibited typical outward rectification (Fig. 2D, red). However, Y308E is toxic (Fig. S2B); therefore, to obtain sufficient channels for electrophysiological characterization, we incubated Y308E-expressing oocytes for 72 h in CBX. These oocytes produced large currents with highly attenuated rectification after CBX removal (Fig. 2D, blue). This attenuation was still visible in oocytes expressing Y308E after only 8 h (Fig. 2D, black). The currents carried by Y308E reversed at a significantly more depolarized potential than those carried by wtPanx1 (Figure 2D inset).

Ion substitution experiments were performed as independent measures of the permeability and selectivity of the Y308E channel. The replacement of Na^+^ by larger cations reportedly does not affect current amplitudes and reversal potentials when Panx1 is stimulated by voltage or deleting the 58 carboxy-terminal amino acids, indicating the absence of cation permeability in the channel (Ma et al., 2012; Romanov et al., 2012; Wang et al., 2014). For Y308E, replacing extracellular Na^+^ with choline^+^ resulted in attenuation of the current amplitudes and a shift of the reversal potential to less negative potentials, indicating choline permeability (Fig. 2E–G). Replacement of Cl^-^ by gluconate^-^ resulted in considerably lower attenuation of the current amplitude for Y308E than that in wtPanx1, demonstrating higher permeability in Y308E (Fig. 2E, wtPanx1 gray). Furthermore, instead of shifting the reversal potential to positive potentials, as observed for wtPanx1, the reversal potential for cells expressing Y308E shifted but remained in the negative domain.

We observed that the phosphomimetic mutation Y308E converted Panx1 into an ATP-release channel, and Y308E as well as Y198 produced currents without activation. We were therefore interested in whether these mutants could be further stimulated by the well-established extracellular activator [K^+^] (Wang and Dahl, 2018). Notably, three mutants, Panx1Y198F/Y308E, Panx1Y198F, and Panx1Y198E, responded to 85 mM K^+^ with increased current amplitudes. However, Y308E-mediated currents did not increase further upon K^+^ application, implying the convergence of K^+^-induced signaling in Y308E (Fig. 2B, Fig. S2C).

Collectively, the results showed that Y308E is in an ATP-releasing, “large” pore conformation of Panx1. We also observed that the phosphomimetic Y308E mutant has different reversal potentials and ion permeabilities than wtPanx1. The permeability of Y308E appears to resemble that of the W74A mutant (Michalski et al., 2020; Ruan et al., 2020).

### Y308E is an ATP-release channel with a dynamic pore and stable wide conformation

Our functional characterization identified Y308E as a spontaneous ATP-release channel permeable to large anions and cations. To gain insight into the “large” ATP-conducting conformation of the Panx1 channel, we obtained a cryo-EM map of Y308E at a resolution of 2.5 Å (Fig. 3F, Fig. S1.1). Although the overall architecture of Y308E is similar to that of other Panx1 structures, Y308 is unique in that it exhibits the widest channel vestibule. In this respect, Y308E is similar to the wider PanxSRC/R75A conformation. Despite the overall high local resolution at the pore (approximately 2.2 Å), the map around W74 remains relatively poor. This is consistent with published Panx1 structures (e.g. (Hussain et al., 2024; Qu et al., 2020; Ruan et al., 2020)). As a result, W74 modeling, and thus the absolute pore dimension, remains ambiguous (Fig. S4A, B, Fig. S5A, B for R75A)(Mim et al., 2021), indicating that W74 is flexible. Subsequently, we performed a focused 3DVA on the ECD of Y308E. Our analysis showed that the densities of R75 and W74 change, which might be correlated; when W74 density diminished, that of R75 pointed away from W74 (Fig. 3A, Movie S5). This movement might create a path for ATP, which was visualized by manually placing an ATP molecule (and its volume filtered to this resolution) on our map (Fig. 3A). Although Y308E is a constitutively active ATP-release channel, it could still be blocked by CBX (Fig. 2C, Fig. S2A), indicating that W74 was accessible for interaction.

**Fig. 3.**
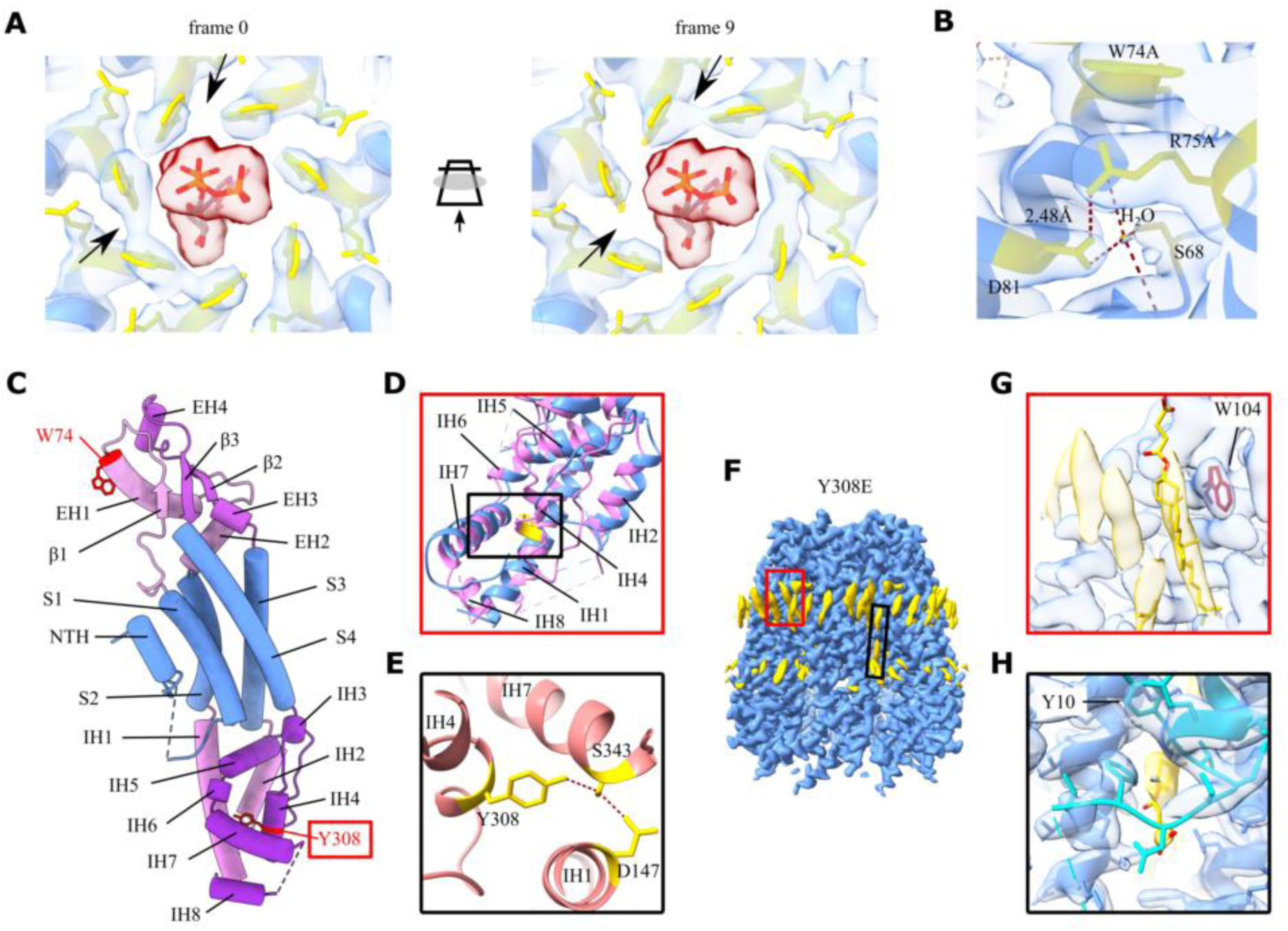
Molecular basis for stabilizing the wide, ATP-releasing conformation. (A**) First (left)** and last (right) volumes of a focused 3DVA volume series for the extracellular domain of Y308E. The animation indicates the viewing direction. Maps are shown in transparent blue, and the Y308E model is in blue. Residues W74 and R75 are highlighted in yellow. The comparison shows that the densities of R75 and W74 change concertedly (indicated by an arrow). A model of ATP (PDB ATP) and its envelope filtered to 3 Å (red) manually placed to visualize the pore. (**B**) Interaction network at the gate connecting W74, R75, D81, and S68 (all yellow) and H_2_O (orange) molecules. The Y308E cryogenic electron microscopy (cryo-EM) map is in transparent blue. Hydrogen bonds and salt bridges are shown (red dashed lines). (**C**) Animation representation and labeling of the model for R74A showing the position of Y308 (red) in the ICD. (**D**) Comparison of the ICDs of Y308E (blue) and R75A (magenta). The position of Y308 on the IH4 helix in the R75A model is highlighted in yellow. The helix is disordered in Y308E. (**E**) Close-up of the hydrogen network (red dashed lines) between residue D147 (IH1), S343 (IH7), and Y308 (IH4) (yellow) in the R75A model (salmon). (**F**) Sharpened cryo-EM map of Y308E at 2.5 Å; the extracellular domain is on top. Non-protein densities (yellow) are positioned in the membrane leaflets. The density penetrating the intra-subunit space has been modeled as two cholesterol hemisuccinate (CHS) molecules. (**G**) CHS on the extracellular side (red box) makes extensive contact with residues in EH2, similar to W104 (red). (**H**) On the intracellular part (black box), the model shows an interaction between CHS and the N-terminus (cyan), stabilizing it in the wide form.

Notably, the geometry of the pore/ECD was stiff, and the distances between the side chains were similar for all structures (Fig. S3). Subsequently, we examined the interaction network of W74 and R75 to understand its influence on R75-W74 dynamics. As reported previously, R75 interacts through a cation-π interaction with W74 and forms a functionally important salt bridge with D81 in an adjacent subunit (Fig. 3B) (Deng et al., 2020; Ruan et al., 2020). In addition, our map showed, for the first time, that a water molecule connects D81 with S68 in the same subunit. This establishes an interaction network connecting the TMD (loop from S1) to the ECD (EH1) and an adjacent subunit (D81). Finally, Y308E appeared restrained into a wide conformation (Movie S7); the largest movements were in the ICD. Collectively, these results suggest that the transition to and maintenance of a wide conformation seem to be essential for ATP release.

### Wide conformation of Y308E is stabilized reorganization of the ICD and the NTH

Y308E was the only mutant with an open Panx1 channel restrained into a wide conformation. To understand this mutation, we examined the environment of Y308 located in the ICD. In our R75A structure, as well as in human wt Panx1, Y308 (Y309 in humans) is part of a helix (IH4). Y308 is in a hydrophobic pocket but forms a hydrogen bond with S343, which forms a hydrogen bond with D147 (Fig. 3E). This triad connects IH1, IH4, and IH7 (Fig. 3D, for nomenclature, see Fig. 3C, Fig. S10). The 3DVA of all our cryo-EM data and Y308E showed that most ICD helices are flexible and likely accessible for modification despite previous assumptions (Ruan et al., 2024). Y198, another phospho-site, is located in IH2 and forms a hydrogen bond with D138, which forms a salt bridge with K202 (Fig. S9A). Phosphorylation of Y198 and/or Y308 modified the geometry of IH1, displacing S2, the helix with the largest displacement in the narrow-to-wide transition (Fig. S8E). In addition, the orientation and composition of helices in the ICD differed between Y308E and R75A, implying a rearrangement of the C-terminus of Panx1.

The second element that likely stabilizes the wide Panx1 conformation is the NTH and the lipids with which it interacts. Although all Panx1 samples are in detergents, we observed non-protein densities reminiscent of residual lipids or detergents (Fig. 3F, yellow). These densities showed a stronger signal at the extracellular interface than at the intracellular interface, similar to the gel-phase lipids observed in connexin hemichannels (Flores et al., 2020). A large density along EH2 connected the hydrophobic core of the membrane with the pore. W104 and F108 formed planar hydrophobic interaction interfaces. We modeled this density as cholesterol hemisuccinate (used during purification) and speculated it constitutes a specific cholesterol-binding site (Fig. 3G). NTH, shown to be an amphipathic helix (Fig. S8D), capped this lipid-filled channel on the pore side of Y308E (Fig. 4H). The hydrophobic residues of this helix are packed between S1 and S4 of the two adjacent subunits (Fig. 3H). This packing may explain the limited mutability of the amino-terminus (Wang and Dahl, 2010). Notably, Y10, a known phosphorylation site, was positioned near the inter-subunit space of Panx1 (Fig. 3H). Changed pY10 interactions might explain why Y10E or Panx1Y10E/Y308E produced undetectable Panx1 currents in oocytes if Y10E channels were expressed in membranes (Fig. 2B). Together, our data on the stabilization of the wide conformation for Y308E suggested that changes occurring during the narrow-to-wide transition are essential for Panx1 function.

### Narrow-wide transition changes interaction interfaces and sequesters the NTH

The W74A functional data imply that replacing W74 with alanine might be insufficient for unstimulated ATP release. However, it seemed that the transition to and maintenance of a wide conformation were essential features of the ATP-releasing conformation. We observed narrow and wide conformations in W74A, R74A, and PanxSRC (movies S2, S4, and S6).

Based on the high-resolution R75A dataset (Fig. 4A, Fig. S1.2), we performed 3DVA on the R75A particle set. We separated particles contributing to the most extreme frames, reconstructed them, and built atomic models (R75A 2.7 Å [wide] vs. 3.4 Å [narrow], Fig. 4 D, Movie S4).

**Fig. 4.**
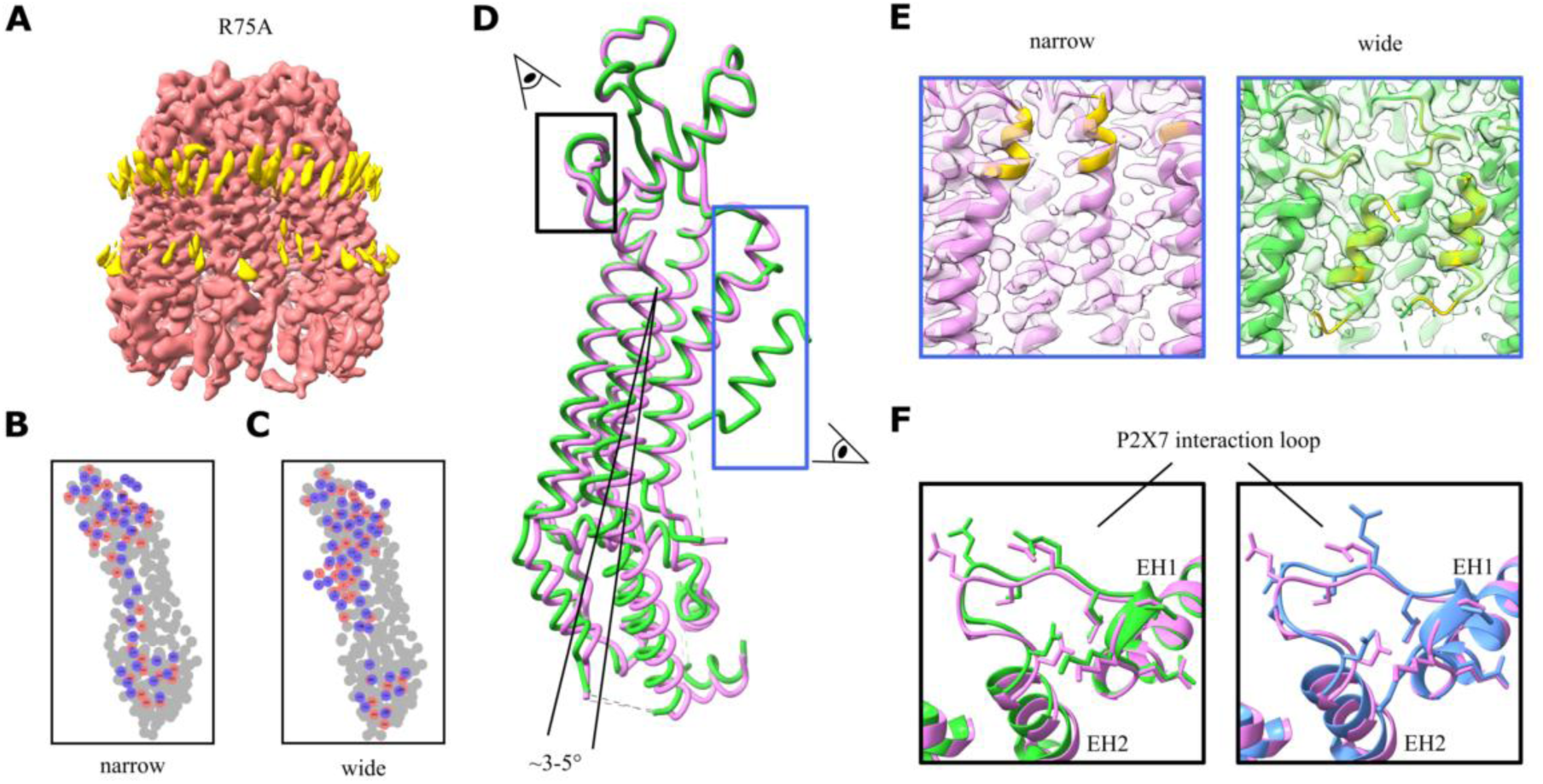
Structural features of the transition from the narrow-to-wide conformation. (**A**) Sharpened cryo-EM map of R75A at 2.8 Å (salmon); the extracellular domain is on top. The non-protein density (yellow) is positioned at the membrane leaflets. (**B**) Residue interaction map for the narrow model between two adjacent R75A protomers. Pink dots represent Chain A, purple dots represent Chain B. (**C**) Residue interaction map for wide conformation as in **B**. (**D**) Single protomer backbone model of the wide (green) and narrow (pink) conformation of R75A. The tilt of the helices in the transmembrane domain (TMD) is between 3° and 5°. The colored boxes highlight areas where the models differ in the transition, and the animated eyes next to the boxes indicate viewing direction in **E** and **F**. (**E**) The two most extreme volume frames from the three-dimensional variability analysis series (map is in transparent gray). (Left) Cryo-EM map and model of the narrow conformation. The C-terminal end of helix S1 (yellow) extends 1–2 turns, and the density for the N-terminal helix (NTH) is not visible nor modeled. (Right) The map and model of the wide conformation and fitted model. Here, the C-terminal helix S1 unwinds (yellow), and NTH (yellow) can be assigned to experimental density (yellow). (**F**) Differences in the P2X7 receptor interacting loop (residues 90–101) (black box in **D**) between the narrow and wide conformations (left, pink vs. green) and between Y308E and the narrow conformation of R75A (right, blue vs. purple). The models are aligned by the extracellular domain.

Overall, the TMD as a whole moved approximately 2.5° away from the pore axis. The S2 helix experienced the largest displacement at approximately 5°, whereas the S3 helix moved approximately 3° (Fig. 4D, Fig. 8D). Consequently, the extracellular helices EH2 and EH1, which contain D81 and W74, respectively, tilted and shifted (Fig. 1F, Fig. 4D). The narrow conformation had tighter transmembrane helix packing at the hydrophobic core of the membrane compared with the wide conformation (Fig. 4B vs. C, Fig. S6A for PanxSRC).

We also observed a non-protein density at and between the Panx1 protomers (Fig. 4A, yellow), which disappeared during the transition from a wide to a narrow conformation (Movie S4). This reduction in the hydrophobic area between subunits has drastic consequences for the amphipathic NTH (Fig. S8D). Like in the probenecid-inhibited Panx1 structure (Kuzuya et al., 2022), the density of the N-terminus in our narrow map was very weak, and the NTH was probably unstructured (Fig. 4E, Fig. S8C for PanxSRC). However, as reported previously, we could not identify lipids in the pore vestibule, nor determine the density in the intracellular domain representing the N-terminus (Kuzuya et al., 2022).

Another difference between the narrow and wide conformations is the C-terminal end of S1. In the narrow map and the probenecid-inhibited Panx1 model, helix S1 contains 1–2 turns that melt into a loop while transitioning into a wider form (Fig. 4E, Movie S4, S8C for PanxSRC, Movie S2). In its wide form, this loop may stabilize the NTH via E57 and connect to residues S68 and R75. Finally, we observed differences in the loop connecting EH1 and EH2 between the two conformations (Fig. 4F, Fig. S8C). This loop mediates the interaction between the ATP-gated P2X7 receptor and Panx1 (Boyce and Swayne, 2017). Thus, opening Panx1 might regulate the formation of the P2X7-Panx1 complex.

## Discussion

Panx1 has “small” chloride-conducting and “large” ATP-permeable states. Proteolytic cleavage of Panx1 has been discussed as an activator of the “large” pore conformation (Narahari et al., 2021; Ruan et al., 2020). However, this hypothesis is problematic because it is an irreversible and pro-apoptotic modification. Furthermore, whether cleaved Panx1 can release ATP shortly after cleavage remains unclear (Narahari et al., 2021; Wang and Dahl, 2018). Alternatively, mechanical stimulation of Panx1 has been shown to increase the conductance and release of ATP (Bao et al., 2004; Reigada et al., 2008). Finally, phosphorylation of Panx1 by SRC tyrosine kinases is a physiological activation mechanism (Lohman et al., 2019). Notably, SRC itself is overexpressed or activated in cancer, ischemia via NMDAR (Weilinger et al., 2012), or vascular dysregulation(Di Salvo et al., 1993). All of these conditions involve Panx1 as well (Laird and Penuela, 2021; Lohman et al., 2019; Molica et al., 2018). Yet, Panx1 phosphorylation has recently been dismissed (Ruan et al., 2024).

Here, we demonstrated that v-SRC increases Panx1-mediated outward currents and conductance. Our mass spectrometry data confirmed phosphorylation at a long-suspected site (Y308) (Weilinger et al., 2016). Weilinger et al. reported in the same study a 10-min delay in the NMDR-SRC-mediated activation of Panx1. This is consistent with our electrophysiological data showing a maximum Panx1 conductance after 10 min (Fig. 1A).

This could be due to the diffusion of v-SRC from the injection site or an indicator that all subunits need to be phosphorylated before Panx1 channels shift into the ‘large’ pore configuration. This interpretation is supported by mass spectrometry that showed that some subunits or whole proteins were not phosphorylated, even after being incubated with v-SRC for over an hour. Therefore, we used phosphomimetic mutants as a condition resembling full phosphorylation. We identified Y308E as the first mutant to create a constitutive ATP-releasing Panx1 channel, reinforcing the hypothesis that phosphorylation of the Y308 residue might be the physiological trigger for ATP release. We hypothesize that different signaling pathways lead to Y308 phosphorylation, which was backed by our observation that external K^+^, a well-established Panx1 activator (Wang et al., 2018), did not further enhance currents in Y308E (Fig. 2C). Notably, Y308E showed currents sooner than wt Panx1, suggesting altered trafficking or a much higher open probability. In contrast, phosphorylation at Y10 or Y150 has been shown to impair trafficking to the cell membrane (Langlois et al., 2023; Nouri-Nejad et al., 2021). Considering that the Y308E pore provides an ATP pathway, we investigated the electrophysiological characteristics of Y308E. Y308E was permeable to large anionic and cationic ions (Fig. 2E). Besides, Y308E attenuated rectification of inward currents (Fig. 2D), similar to the alanine mutant, in which the pore-constricting residue W74 was substituted (Michalski et al., 2020; Ruan et al., 2020). However, all Y308E actions were still blocked by CBX (Fig. S2A), indicating that the W74 of the phosphomimetic was still accessible. Phosphorylation at Y198, another phosphorylation site, seems somewhat redundant when Y308 is phosphorylated, implied by the Y198F/Y308E double mutant that still releases ATP (Fig. 2A). Y198 may not be as relevant because Y198 has been proposed to be constitutively active (DeLalio et al., 2019) and thus be ruled out as a functional switch.

We gained structural insight into ATP release by solving the structures of dephosphorylated and v-SRC-phosphorylated wtPanx1 (PanxSRC), the pore-lining mutants W74A and R75A, and the ATP-releasing mutant Y308E. All structures showed a typical Panx1 architecture (as reviewed in (Syrjanen et al., 2021)). To the best of our knowledge, this study is the first to examine the flexibility of Panx1 domains as a functionally important factor. First, we investigated W74, which was assumed to be a key residue for ATP release (Michalski et al., 2020; Qu et al., 2020; Ruan et al., 2020). Not only was the density around W74 unusually weak (Fig. S4, Fig. S5), but the highest resolution map (Y308E) demonstrated that the pore residues W74 and R75 were flexible. This flexibility may also explain why the electrophysiological properties of Y308E are similar to those of W74A (or R75A). The flexibility of W74 may allow ATP to pass through the pore in Y308E. The flexibility also implies that W74 modeling in most structures is ambiguous at best (shown in (Mim et al., 2021)) and casts doubt on whether these residues are static. However, interaction with ATP or CBX in the pore may reduce flexibility and explain the inhibitory effect at high concentrations (Ruan et al., 2020). The Y308E model suggested that the R75 interaction network (D81 and S68) pulled R75 away from W74. Our results question the established role of W74 in ATP permeation, given that our map showed that W74A generates a pore wide enough (approximately 12 Å) for the passage of ATP and evidence that W74A is not an ATP-releasing channel (Qiu et al., 2012).

We speculated that the conformational changes of Panx1 as a whole may explain ATP release. We observed that W74A, R75A, and PanxSRC altered between the two states; we named these narrow and wide (Movies S2, S4, and S6). The exception is Y308E, which resides in a wide state with little movement in the TMD (Movie S7). We speculate that ATP-releasing Panx1 has to be in a wide conformation because, in the narrow conformation, the NTH is most likely disordered and located in the vestibule. The detached NTH created a negatively charged diffusion barrier for ATP in the narrow conformation. A recent study on the NTH in connexins showed that its NTH is flexible and interacts with large molecules permeating the channel, supporting our model (Gaete et al., 2024). The NTH of Panx1 may also be necessary to prevent lipids from entering the channel, as suggested previously (Kuzuya et al., 2022), particularly in the wide conformation; however, we did not observe any lipid density inside the pore vestibule. We hypothesize that the wide conformation of Y308E was maintained by electrostatic repulsion in the presence of D147 and the relatively hydrophobic environment around Y308. This is in contrast with the environment around Y198 with two lysine residues (K202 and K345 in another subunit) that could interact with a phosphorylated Y198. This may explain why the phosphorylation of Y308 is more impactful. The ICD was dynamic. As such, Y198 and Y308 were most likely accessible for modification, contrary to earlier speculations (Ruan et al., 2024). ICD movements may also steer the C-terminal region. As in most published Panx1 structures, the C-terminus was not resolved in our studies. However, functional data strongly suggest that the C-terminus dives deep into the channel pore, which is also supported by the single publication resolving the C-terminus (Zhang et al., 2021). Therefore, no conclusions can be made about the contribution of the C-terminus to Panx1 channel activity which has been shown to be associated with the extracellular pore and is important for permeability. Finally, we also observed the conformation-dependent movement of a P2X_7_ receptor binding motif (Boyce and Swayne, 2017) in Panx1. While P2X_7_ receptor-dependent activation of Panx1 is thought to be driven by SRC (Iglesias et al., 2008), our model offers an additional regulatory element.

In conclusion, we propose a “selective rigidity” gating mechanism for ATP release (Fig. 5). Although this study did not address it explicitly, it offers a model for the mechanosensitive activation of Panx1 (Locovei et al., 2006a; Reigada et al., 2008). A compression or relaxation of transmembrane helices, similar to the narrow vs. wide conformation in Panx1, has been observed in other mechanosensitive channels (Deng et al., 2020; Mount et al., 2022). Certain membrane curvatures, or lipids, may bias the conformational equilibrium. However, further studies of Panx1 in different membrane systems are necessary to validate this idea.

**Fig. 5.**
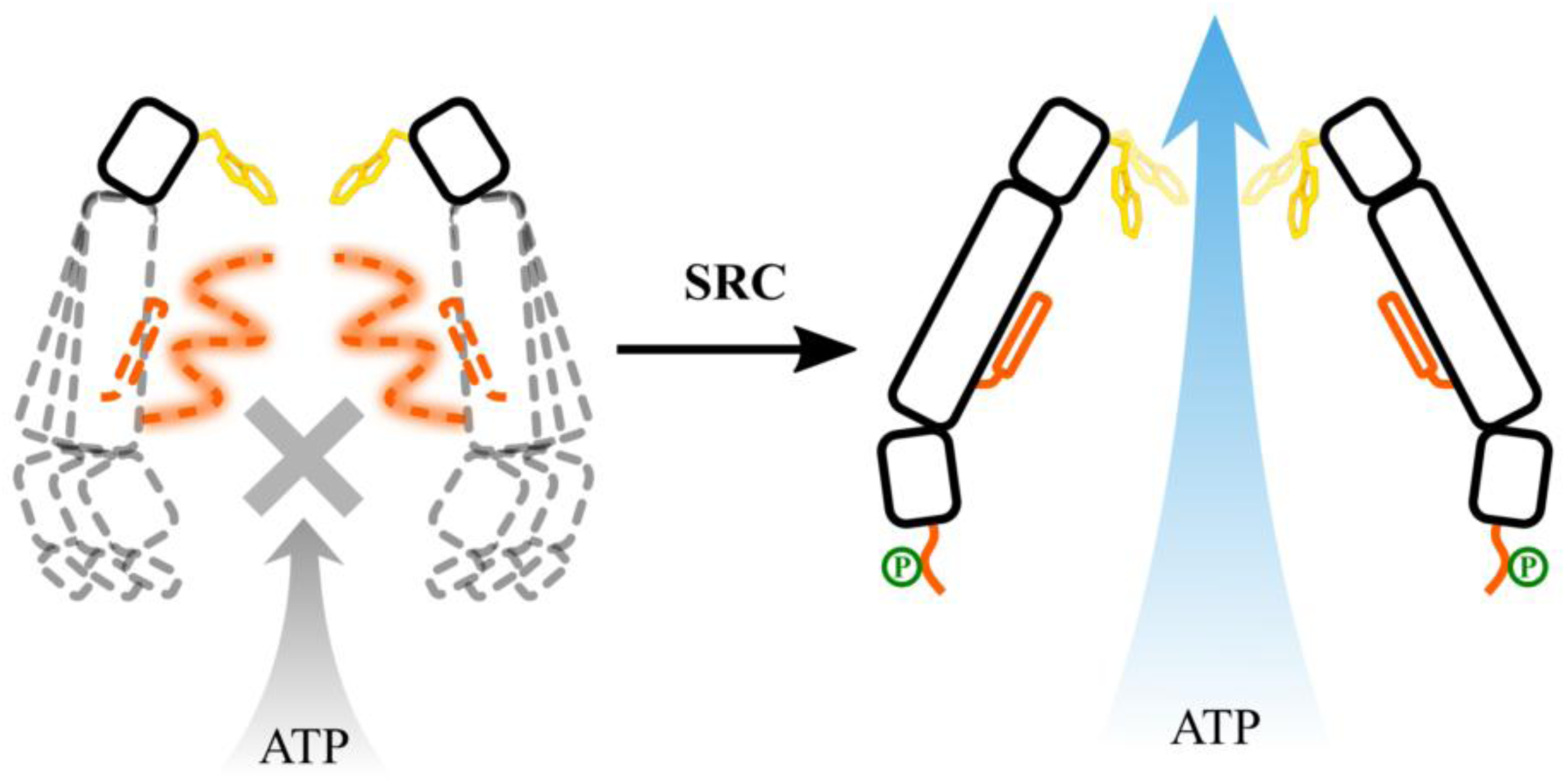
“Selective rigidity” mechanism for ATP release by Panx1. (Left) In a non- or partially phosphorylated state, the transmembrane domains and intracellular domains are flexible. Panx1 transitions between a narrow and wide state. Consequently, the N-terminal helix is only partially ordered (light orange) or disordered (dashed orange). The residue W74 (yellow) at the extracellular gate shows some degree of flexibility. (Right) Phosphorylation (green) at residue Y308 causes the intracellular helix (H1) and transmembrane helix (S2) to tilt away. This state is stabilized and maintained. As a result, the NTH is formed and sequestered away (orange). The S2 movement takes other helices and destabilizes the R75-W74 interactions, allowing W74 to move more freely (yellow and blurred). As a result, ATP can pass unimpeded through the vestibule and flexible pores.

Nevertheless, our integrative approach offers the first model showing how phosphorylation changes the dynamics and permeability in Panx1, which may be a common phenomenon in large-pore channels. This mechanism offers an avenue for the development of conformation modulators, like probenecid, for diseases from cancer to chronic pain.

## STAR Methods

### Constructs and cloning

The full-length mouse Panx1 subcloned into the pFastBac1 vector, a gift from Daniela Boassa/Gina Sosinsky, contained a Tobacco Etch Virus (TEV) protease recognition sequence (Glu-Asn-Leu-Tyr-Phe-Gln-Gly) and 6×His tag at the C-terminus. A Phusion Site-directed Mutagenesis Kit (Life Technologies, Carlsbad, CA, USA) was used to generate all point mutations (W74A, R75A, and Y308E). All primers were synthesized using integrated DNA technologies. Positive clones were verified using Sanger sequencing by Eurofins Genomics (Luxembourg) and Microsynth AG (Balgach, Switzerland). All correct pFastBac1 constructs were transformed into DH10 Multibac *E. coli* (Geneva Biotech, Geneva, Switzerland) to generate recombinant Bacmid constructs using the Bac-to-Bac^TM^ Baculovirus Expression System (Life Technologies) according to the manufacturer’s instructions. Bacmids from positive white colonies were extracted using ethanol precipitation for viral production.

### Preparation of oocytes

Ovaries were harvested from adult female African clawed frogs (*Xenopus laevis*), cut into small pieces, and incubated in collagenase (2.5 mg/mL; Worthington Biochemical Corporation, Lakewood, NJ, USA) in calcium-free oocyte Ringer’s (OR) solution with stirring at one turn per second at room temperature (23 °C). Typically, the incubation period for oocytes to be separated from follicle cells is 3 h. After thorough washing with regular OR (82.5 mM NaCl, 2.5 mM KCl, 1 mM Na_2_HPO_4_, 1 mM MgCl_2_, 1 mM CaCl_2_, and 5 mM HEPES), oocytes devoid of follicle cells and with uniform pigmentation were selected and stored in OR at 18 °C for 18 h to 3 days before electrophysiological analysis at room temperature.

### Preparation of mRNA and electrophysiology

Plasmids containing mouse Panx1 or its mutants in pCS2 were linearized with NotI. *In vitro* transcription was performed with SP6 polymerase using the mMESSAGE mMACHINE™ T7 Transcription Kit (Ambion, Austin, TX, USA). mRNA was quantified using absorbance (260 nm), and the proportion of full-length transcripts was determined by agarose gel electrophoresis. *In vitro*-transcribed mRNAs (60 nL at approximately 1 µg/µL) were injected into *X. laevis* oocytes. Cells were maintained in regular Ringer’s solution with streptomycin (10 mg/mL). Whole-cell membrane currents of the oocytes were measured using a TEVC (Gene Clamp 500 B; Axon Instruments, San Jose, CA, USA) under constant perfusion according to the protocol described in Fig. 2. The glass pipettes were pulled using a P-97 Flaming/Brown puller (Sutter Instrument Company, Novato, CA, USA). Two electrophysiological protocols were used: to determine membrane conductance, the membrane potential was held at –60 mV, and small test pulses lasting 6 s at –48 mV, or test pulses from –60 to +60 mV, were applied at a rate of 0.1 Hz; alternatively, voltage ramps lasting 35 s were applied, typically from –100 to +100 mV.

Because of the shortened lifetime of oocytes expressing Panx1Y308E, only a short time window of 4–8 h after mRNA injection was available for electrical recording and determination of ATP release. wtPanx1 and all other mutants were analyzed 24–48 h after mRNA injection.

### ATP-release assay

ATP flux was determined using luminometry. Briefly, oocytes were analyzed 8 h after injecting Panx1Y308E mRNA or 2 days after injecting wtPanx1. Next, 90 µL of oocyte supernatant were added to 40 µL of luciferase-luciferin solution (Promega, Madison, WI, USA) to assay luciferase activity. Data from oocytes with visible damage were excluded from the analysis.

### Transfection and virus production

The Sf9 cells used in this study were purchased from Life Technologies. The cells were routinely maintained in our laboratory, not authenticated, and tested negative for *Mycoplasma* contamination. The Sf9 cells were grown in suspension at 28 °C at 110 rpm on a shaker in sf-900^TM^ III medium (Invitrogen, Carlsbad, CA, USA) in Corning^®^ Erlenmeyer shake flasks (non-baffled with vent cap; Corning Life Sciences, Corning, NY, USA).

Adherent Sf9 cells were transfected with Bacmid constructs using X-tremeGENE™ 9 DNA Transfection Reagent (Life Technologies) following the manufacturer’s protocol and Bac-to-Bac^TM^ Baculovirus Expression System user guide (Life Technologies) to produce the virus. Brilliant Blue FCF (10 µM) was added as an inhibitor during transfection and virus production only for the Y308E mutant. Viruses were kept at 4 °C and protected from light until use.

### Protein expression and purification

A test expression was performed to determine the optimal virus-to-cell ratio and harvest time. Suspended Sf9 cells at a density of 2 × 10^6^ cells/mL were infected at different virus-to-cell ratios. Equal amounts of cells were harvested by centrifugation 2, 3, 4, and 5 days post-infection. Western blotting was performed using an anti-His antibody (AD1.1.10; NB100-64768; Novus Biologicals, Centennial, CO, USA) to determine the conditions yielding the highest expression level.

For high Panx1 expression, a suspension of Sf9 cells at a density of 2 × 10^6^ cells/mL was infected with the optimal virus amount determined using test expression. Brilliant Blue FCF (10 µM) was added as an inhibitor only for Y308E mutants. Cells were pelleted based on test expression results (approximately 4 days after infection) and stored at −80 °C until further use.

Infected Sf9 cells (pellets from 1 L of culture) were thawed on ice and resuspended in lysis buffer (150 mL NaCl, 50 mM HEPES at pH 8.0, and COmplete™ Protease Inhibitor; Roche, Basel, Switzerland). Thereafter, cells were lyzed by sonication at 65% power for 8 min with 5 s on and 25 s off. Cell debris was removed by centrifugation at 4,000 ×*g* for 20 min at 4 °C. Membranes from the supernatant were harvested by ultracentrifugation (Ti70.1 rotor; Beckman Coulter, Brea, CA, USA) at 105000 g for 1 h. Membrane pellets were homogenized using a Dounce Tissue Grinder (DWK Life Sciences, Malvern, PA, USA) and solubilized with 2% *N*-dodecyl-ß-D-maltoside (DDM) and 0.4% cholesterol hemisuccinate (CHS; Sigma-Aldrich, St. Louis, MO, USA) in solubilization buffer (200 mM NaCl, 50 mM HEPES at pH 8.0, with cOmplete™ protease inhibitor) for 2 h at room temperature. The insoluble material was removed by ultracentrifugation (Ti70.1 rotor; Beckman Coulter) at 105000g for 1 h. Subsequently, the supernatant was adjusted to 20 mM imidazole and bound to 2 mL (bed volume) Ni-NTA (G-Bioscience, St. Louis, MO, USA) using a batch protocol for 2 h at 4 °C. Subsequently, the resin was washed with a 20× bed volume of washing buffer 1 (150 mM NaCl, 50 mM HEPES at pH 8.0, 0.1% DDM [Avanti Polar Lipids, Alabaster, AL, USA], 0.02% CHS, and 20 mM imidazole), followed by washing buffer 2 (150 mM NaCl, 50 mM HEPES at pH 8.0, 0.02% glycol-diosgenin [GDN, Avanti Polar Lipids], and 20 mM imidazole) as detergent exchange. Afterward, proteins were eluted with elution buffer (150 mM NaCl, 50 mM HEPES at pH 8.0, 0.02% GDN, and 250 mM imidazole) and concentrated using a 100-kDa cutoff concentrator (Sartorius, Göttingen, Germany). The concentrated protein was further purified by size-exclusion chromatography (SEC) using a Superose 6 Increase 10/300 GL column (Cytiva, Uppsala, Sweden) with cell lysis buffer containing 0.02% GDN. Peak fractions were concentrated using a 100-kDa cutoff concentrator (Sartorius) for cryo-EM sample preparation. Sodium dodecyl-sulfate polyacrylamide gel electrophoresis and western blotting were performed to confirm expression and purity.

### Phosphorylation and de-phosphorylation

After eluting Panx1 from Ni-NTA, the buffer was exchanged to 150 mM NaCl, 50 mM HEPES at pH 8.0, 0.02% GDN, and dephosphorylated with 26 ku of Lambda Phosphatase (Sigma-Aldrich) at 4 °C overnight. The next morning, the reaction was separated using SEC, as described above. Thereafter, the fractions were pooled and concentrated to 1.6 mg/mL. Next, 30 µL were adjusted using 150 mM NaCl, 50 mM HEPES at pH 8.0, 0.02% GDN, 5 mM MnCl_2_, 5 mM Mg_2_Cl, 0.25 mM ATP, 6 µM NaVO_4_, 2 µL v-SRC (Sigma-Aldrich) for 1–2 h before cryo-EM sample preparation. The remaining samples were frozen and stored for mass spectrometry.

### Cryo-EM sample preparation and data collection

All samples were vitrified in a VitroBot Mark IV (Thermo Fisher Scientific, Waltham, MA, USA) at 6 °C and 100% relative humidity. Glow-discharged QuantiFoil grids (MicroTools, Simsbury, CT, USA) were used for all samples. Specific blotting conditions and protein concentrations differed between samples. Fluorinated Fos-Choline 8 (FFC8) was added. The concentration, grid type, and glow discharge conditions were as follows: Panx1SRC 1.6 mg/mL, QF Cu300 1.2/1.3, 20 mA/60 s; Panx1Dephos 1.3 mg/mL, QF Cu300 1.2/1.3, 20 mA/60 s; R75A 6 mg/mL, QF AUtra300, 1.2/1.3, 12 mg/mL, 5 mM FFC8, QF Cu300, 1.2/1.3, 40 mA/60 s; W74A 4 mg/mL, 5 mM FFC8, QF Cu300, 1.2/1.3; Y308E 3.5 mg/mL, 5 mM FFC8, QF Cu300, 1.2/1.3, 40 mA/60 s.

Cryo-EM data were collected on a 300-kV Thermo Fisher Titan Krios instrument (Thermo Fisher Scientific) equipped with a Gatan BioQuantum energy filter (Gatan Inc., Westlake Village, CA, USA) and either a Gatan K3 (Gatan Inc.) or Falcon4 Direct Electron Detector (Thermo Fisher Scientific). The nominal magnifications of each sample are presented in Table 1. The total dose was between 40 and 50 e^−^/Å^2^. The defocus was varied between –0.6 and –2.8 Å, depending on the sample. Data were collected using the EPU software (Thermo Fisher Scientific).

**Table 1.** Cryogenic electron microscopy and model statistics.

|  | Panx1S<br>RC | Panx1<br>SRC<br>wide | Panx1<br>SRC<br>narrow | Panx1Y3<br>08E | Panx1R7<br>5A | Panx1<br>R75A<br>wide | Panx1<br>R75A<br>narrow | Panx1W7<br>4A | Panx1D<br>eph |
| --- | --- | --- | --- | --- | --- | --- | --- | --- | --- |
| Magnification | 165000 | 165000 | 165000 | 130,000 | 105,000 | 105,000 | 105,000 | 165000 | 105000 |
| Voltage<br>(kV) | 300 | 300 | 300 | 300 | 300 | 300 | 300 | 300 | 300 |
| Electron<br>exposure<br>(e-/Å<sup>2</sup>) | 50 | 50 | 50 | 50 | 50 | 50 | 50 | 50 | 50 |
| Defocus<br>range (µm) | 1.0–2.5 | 1.0–2.5 | 1.0–2.5 | 0.6–1.8 | 0.9–2.1 | 0.9–2.1 | 0.9–2.1 | 1.0–2.5 | 1.0–2.0 |
| Pixel size<br>(Å) | 0.704Å | 0.704Å | 0.704Å | 0.65Å | 0.825Å | 0.825Å | 0.825Å | 0.704Å | 0.86Å |
| Symmetry<br>imposed | C7 | C7 | C7 | C7 | C7 | C7 | C7 | C7 | C7 |
| Initial<br>particle<br>images<br>(no.) | 745,409 |  |  | 957,362 | 1,660,043 |  |  | 931,931 | 1,590,024 |
| Final<br>particle<br>images<br>(no.) | 112,178 | 27,144 | 39,410 | 168,845 | 160,375 | 24,631 | 28,395 | 71,613 | 94,273 |
| Map<br>resolution<br>(Å) | 3.56Å | 3.31Å | 3.93Å | 2.45Å | 2.86Å | 2.74Å | 3.52Å | 3.75Å | 4.68Å |
| FSC curve | normal | auto-tightening | normal | auto-tightening | auto-tightening | auto-tightening | normal | auto-tightening | normal |
| FSC<br>threshold | 0.143 | 0.143 | 0.143 | 0.143 | 0.143 | 0.143 | 0.143 | 0.143 | 0.143 |
| Map<br>resolution<br>range (Å) | 1.5–48.7 | not determined | not determined | 2.2–22.8 | 1.8–34.2 | not determined | not determined | 3.4–7.9 | 1.8–6.7 |
| Initial | Q9JIP4 from AlphaFold-DB |  |  |  |  |  |  |  |  |
| model used<br>(PDB ID) |  |  |  |  |  |  |  |  |  |
| Model<br>resolution<br>(Å) | 3.43 | 3.33 | 4.15 | 2.43 | 2.83 | 2.73 | 3.53 | 3.83 | 5.58 |
| FSC<br>threshold | 0.143 | 0.143 | 0.143 | 0.143 | 0.143 | 0.143 | 0.143 | 0.143 | 0.143 |
| Map<br>sharpening<br>B factor<br>(Å<sup>2</sup>) | not applicable, refinement against an unsharpened map |  |  |  |  |  |  |  |  |
| Model<br>compositio<br>n |  |  |  |  |  |  |  |  |  |
| Non-<br>hydrogen<br>atoms | 15911 | 15043 | 14819 | 17990 | 17934 | 17934 | 17038 | 17913 | 17976 |
| Protein<br>residues | 1960 | 1855 | 1827 | 2177 | 2226 | 2226 | 2114 | 2226 | 2226 |
| Ligands | 0 | 0 | 0 | 14 | 0 | 0 | 0 | 0 | 0 |
| Water<br>molecules | 0 | 0 | 0 | 21 | 0 | 0 | 0 | 0 | 0 |
| R.m.s.<br>deviations |  |  |  |  |  |  |  |  |  |
| Bond<br>lengths (Å) | 0.009 | 0.003 | 0.006 | 0.003 | 0.002 | 0.003 | 0.006 | 0.003 | 0.005 |
| Bond<br>angles (°) | 1.039 | 0.687 | 1.037 | 0.647 | 0.584 | 0.587 | 0.766 | 0.659 | 0.968 |
| Validation |  |  |  |  |  |  |  |  |  |
| Clashscore | 10.41 | 2.94 | 18.72 | 0.82 | 0.85 | 1.32 | 2.97 | 5.40 | 14.27 |
| Poor<br>rotamers<br>(%) | 0.00 | 0.30 | 3.50 | 0.05 | 0.00 | 0.05 | 0.62 | 1.08 | 1.37 |
| Ramachan<br>dran plot |  |  |  |  |  |  |  |  |  |
| Favored<br>(%) | 98.12 | 97.20 | 95.16 | 96.84 | 95.94 | 95.44 | 95.27 | 96.18 | 96.82 |
| Allowed<br>(%) | 1.88 | 2.75 | 4.55 | 3.16 | 3.96 | 4.19 | 4.39 | 3.73 | 2.67 |
| Disallowed<br>(%) | 0.00 | 0.06 | 0.28 | 0.00 | 0.09 | 0.37 | 0.34 | 0.09 | 0.51 |

### Cryo-EM single particle analysis

All data were processed using CryoSparc version 4.1-4.5 (Structura Biotechnology Inc., Toronto, ON, Canada) (Punjani et al., 2017). The raw movies were motion-corrected using patch motion correction software. The contrast transfer function (CTF) was estimated using the software’s patch CTF estimation. The micrographs were curated based on the CTF fit resolution, motion distance, and ice quality. The particles were manually selected from a small subset of images and used for Topaz training (Bepler et al., 2020). After training and picking, the particles from all movies were curated using two-dimensional classification. The remaining particles were further sorted using *Ab Initio*/Heterogeneous refinement cycles. The Y308E and R75A datasets contained a small number of Panx1 dimers (10–20%), which were removed using *Ab Initio*/Heterogeneous refinement cycles. We also combined the two datasets for R75A because the structures and 3DVA were similar for both particle sets. The final particles for each sample were subjected to Non-Uniform Refinement with or without C7 symmetry (Punjani et al., 2020). These particles underwent optimization using a reference-based local motion correction and subsequent refinement.

### 3DVA and refinement

For CryoSparc 3DVA (Punjani and Fleet, 2021), the refined particles expanded symmetrically (C7). The focusing mask was based on a model we created. 3DVA was performed on either the ECD or the whole protein with a resolution cutoff of 2.5–3 Å. Before reference-based motion correction and extraction of the final box size, an initial 3DVA was performed and used only for illustrative purposes. After refinement using motion-corrected particles, we conducted the final 3DVA. To separate the particles associated with individual volumes in a series, we recorded the particles for ten “intermediate” frames. We used only the first and last frames representing the most different volumes. Symmetry-related duplicates were removed, and the remaining particles were subjected to local refinement using C7 symmetry.

### Model building, validation, and structure analysis

The AlphaFold prediction of the mouse Panx1 protomer (UniProt accession code Q9JIP4) was downloaded from AlphaFold-DB (DeepMind, London, UK) (Varadi et al., 2022) and rigidly fitted into the cryo-EM map, along with six additional copies generated by applying the C7 symmetry of the map using ChimeraX version 1.6.1 (Pettersen et al., 2021). Segments of the atomic model that were not supported by any density were deleted, and point mutations were applied to match the corresponding Panx1 mutants that produced each cryo-EM map. The resulting models were refined against the cryo-EM map by interactive molecular dynamics flexible fitting (iMDFF) using ISOLDE version 1.6.0 (Croll, 2018) within ChimeraX. A post-processed map generated by deepEMhancer (Sanchez-Garcia et al., 2021) using tightTarget weights was used as a visual aid for interactive model refinement in ISOLDE (but was not used to drive MDFF) following the strategy outlined in (Sanchez-Garcia et al., 2024). Each residue in the first protomer (chain A) was visually inspected at least once. The resulting models were submitted for a final real-space refinement using phenix.real_space_refine from the Phenix Suite version 1.20.1-4487 (Afonine et al., 2018b) using the settings file generated by ISOLDE. The final atomic models were validated against the corresponding cryo-EM maps using phenix.validation_cryoem (Afonine et al., 2018a). All images were generated using the ChimeraX or Chimera software.

### Mass spectrometry

Dephosphorylated and v-SRC-phosphorylated mPanx1 were analyzed using untargeted bottom-up proteomics. For each sample, the volume was adjusted to 100 µL (50 mM HEPES). Next, samples were denatured at 60 °C for 1 h, followed by reduction with DTT at 95 °C for 30 min and alkylation with chloroacetamide at room temperature for 20 min at end concentrations of 4 mM. Chloroacetamide was blocked by adding DTT. Trypsin (trypsin sequencing grade modified; Pierce, Thermo Fisher Scientific) was dissolved (10% ACN, 50 mM HEPES), added at a 1:50 (w/w) ratio, and digestion was performed at 37 °C overnight. Each sample was separated for 1 h using a Dionex Nano LC-System (Thermo Fisher Scientific) with a 5–40% ACN gradient coupled with a QExactive HF Mass Spectrometer (Thermo Fisher Scientific) operating in DDA mode. The PEAKS Studio software (Bioinformatics Solutions Inc., Vancouver, BC, Canada) was used to search the *Mus Musculus* and *Spodoptera Frugiperda* Uniprot databases for protein and peptide identification, limited to a false discovery rate of 1%. The search results were filtered to contain only the data points from all three replicates.

### Ethic statements

All procedures were approved by the University of Miami Institutional Animal Care and Use Committee and conducted in accordance with the Guiding Principles for Research Involving Animals and Human Beings of the American Physiological Society. The study details were in accordance with the ARRIVE guidelines for the use of laboratory animals.

## Resource availability

### Materials availability

Plasmids generated in this study are available upon request.

## Acknowledgments

Data were collected at the Cryo-EM Swedish National Facility funded by Knut and Alice Wallenberg, Family Erling Persson, Kempe Foundations, SciLifeLab, Stockholm University, and Umeå University.

## Author contributions

Conceptualization: CM, GD, RBS; Methodology: CM, QZ, GD, RBS, GG; Investigation: CM, JW, GG; Visualization: QZ, CM, GD, RBS, GG; Funding acquisition: CM; Project administration: CM, RBS, GD; Supervision: CM, CB, GD, RBS; Writing – CM, GD, QZ, GG, DK, RBS; Writing – review & editing: CM, GD, QZ, GG, DK, CB, RBS.

